# Brain connectivity measures improve modeling of functional outcome after acute ischemic stroke

**DOI:** 10.1101/590497

**Authors:** Sofia Ira Ktena, Markus D. Schirmer, Mark R. Etherton, Anne-Katrin Giese, Carissa Tuozzo, Brittany B Mills, Daniel Rueckert, Ona Wu, Natalia S. Rost

## Abstract

**Background:** The ability to model long-term functional outcomes after acute ischemic stroke (AIS) represents a major clinical challenge. One approach to potentially improve prediction modeling involves the analysis of connectomics. The field of connectomics represents the brain’s connectivity as a graph, whose topological properties have helped uncover underlying mechanisms of brain function in health and disease. Specifically, we assessed the impact of stroke lesions on rich club (RC) organization, a high capacity backbone system of brain function.

**Methods:** In a hospital-based cohort of 41 AIS patients, we investigated the effect of acute infarcts on the brain’s pre-stroke RC backbone and post-stroke functional connectomes with respect to post-stroke outcome. Functional connectomes were created utilizing three anatomical atlases and characteristic path-length (*L*) was calculated for each connectome. The number of RC regions (N_RC_) affected were manually determined using each patient’s diffusion weighted image (DWI). We investigated differences in *L* with respect to outcome (modified Rankin Scale score (mRS); 90-days; poor: mRS>2) and the National Institutes of Health Stroke Scale (NIHSS; early: 2-5 days; late: 90-day follow-up). Furthermore, we assessed the effect of including N_RC_ and *L* in ‘outcome’ models, using linear regression and assessing the explained variance (R^2^).

**Results:** Of 41 patients (mean age (range): 70 (45-89) years), 61% were male. There were differences in *L* between patients with good and poor outcome (mRS). Including NRC in the backward selection models of outcome, R^2^ increased between 1.3- and 2.6-fold beyond that of traditional markers (age and acute lesion volume) for NIHSS and mRS.

**Conclusion:** In this proof-of-concept study, we showed that information on network topology can be leveraged to improve modeling of post-stroke functional outcome. Future studies are warranted to validate this approach in larger prospective studies of outcome prediction in stroke.

## Introduction

Stroke is a leading cause of long-term adult disability^1^ with significant public health burden^2^. Importantly, the ability to individually prognosticate stroke outcomes in the acute setting remains challenging^3^, due to the complex mechanisms of post-stroke recovery and the multitude of clinical and radiographic variables that differentially affect patient outcomes^4–6^.

Magnetic resonance imaging (MRI) allows the mapping of anatomical regions and the pathways of their interconnections through diffusion weighted imaging (DWI) or functional co-activation through functional MRI (task-based or resting state (rsfMRI)), producing a comprehensive description of the brain’s structural and/or functional connectivity. Connectomics involves conceptualizing the brain as a graph and allows the exploration of topological properties of brain connectivity with network theoretical measures^7^. This has led to fundamental insights into the brain’s organization^8–12^, resilience to injury^13–15^, and alterations due to disease^16–20^. Associations between structural features, such as white matter microstructural integrity, and functional post-stroke outcome have recently been established^21^. However, the effect of premorbid structural and/or functional brain connectivity organization on recovery after stroke and its role in resilience to damage is yet to be fully elucidated.

A so-called rich club (RC) organization has been described in the human connectome^8,10^, comprising a set of regions which are thought to form an information backbone, crucial for brain function, and susceptible to disease^22^. Van den Heuvel and Sporns^8^ identified six bilateral regions belonging to the RC, three cortical (precuneus, superior frontal and parietal cortex) and three subcortical regions (putamen, hippocampus and thalamus), where RC regions are hubs that mediate long-distance connections between brain modules^23^. This demonstrated their critical role for information integration, adaptive behavior^24^ and cognitive tasks^25^. Targeted attacks on their connections can have a significant impact on global network efficiency^8^ and have been shown to lead to functional deficits in disorders like Alzheimer’s disease^26^. Importantly, stroke location has been identified as an independent determinant of cognitive outcome^27,28^, in addition to widely accepted clinical factors, such as age^29^ and stroke lesion size^30,31^. Furthermore, a strong coupling between brain hubs, especially those lying in the cerebral cortex, and regional blood flow has been unveiled during rest as well as in response to task demands^32^. Consequently, the investigation of damage to RC regions in stroke patients is intriguing. The effect of focal injury caused by stroke on large-scale brain networks has been recently explored along with network alterations in brain tumor and traumatic brain injury^33^, however, without a clear mapping between the anatomical lesion site and its topological characteristics within the brain network. Functional connectivity has been previously explored in longitudinal studies of motor recovery after stroke^34,35^ and significant correlations between interhemispheric resting-state connections and functional performance have been identified^36,37^. Nevertheless, the effect of focal ischemic stroke lesions on whole-brain functional organization estimated before and after stroke have not been investigated.

In this study, we examine the functional network organization in AIS patients and the lesion location in relation to network topography with respect to functional outcome. Here, we assessed the impact of ischemic insults on brain regions that constitute the RC backbone, as well as functional network topology at a global level on brain recovery, in a prospective, hospital-based cohort. We hypothesize that models incorporating connectivity information, specifically the characteristic path length in the acute phase and the number of RC regions affected by the stroke lesion, will improve the prediction of a patient’s functional outcome. Using multivariate linear regression, we conclude that the connectivity metrics obtained early in the course of acute ischemic stroke can be used to better understand the mechanisms underlying variability in post-stroke functional outcomes.

## Materials and Methods

### Patient population

AIS patients were enrolled in the SALVO (Statins augment small vessel function and improve stroke outcomes) study after admission to the Emergency Department at Massachusetts General Hospital. The study was approved by the Institutional Review Board and all participants, or their surrogates, gave written informed consent at the time of enrolment. AIS was defined as: (a) acute onset of focal neurological symptoms consistent with cerebrovascular syndrome, (b) MRI findings consistent with acute cerebral ischemia, and (c) no evidence of other neurological disorders to explain the symptoms. Subjects with moderate to severe white matter hyperintensity (WMH) burden defined as Fazekas^38^ grade ≥2 in any of the three categories (periventricular, deep lesion extent and deep lesion count) were eligible for enrolment in this study. Participants with medical contraindications to gadolinium-based contrast agents were excluded from this study.

### Clinical assessment

Upon admission to the hospital, the National Institutes of Health Stroke Scale score^39^ (NIHSS; 0 (no symptoms) - 42) was recorded for each patient by a trained neurologist^40^. Utilizing NIHSS as a pseudo outcome score, post-stroke functional outcome was assessed during two follow-up assessments: (1) within 2-5 days after admission (average 2.6 days, “early” in-hospital follow-up) and (2) at 90 days (“late” follow-up). Additionally, the modified Rankin Scale score^41^ (mRS; 0 (no symptoms) - 6 (death)) was recorded at “late” follow-up to assess functional status of the patients. mRS focuses on the assessment of functional independence (ability to return to independent living, including ambulation without assistance), and is widely used in stroke clinical trials based on its high utility and reliability^42–44^. In this study, mRS ≤2 was considered ‘good’ outcome (minor disability but patient is functionally independent), while mRS>2 was considered ‘poor’ outcome (significant disability, loss of functional independence, including death).

### Data acquisition

Patients enrolled in the SALVO study underwent a research protocol MRI, including structural, diffusion and functional imaging, in the hospital at 2-5 days after admission. A T1-weighted image was acquired with the following parameters: in-plane resolution, 0.430 mm; slice thickness, 6mm; matrix size, 480×512; number of slices: 28. Gradient-echo echoplanar imaging (EPI) data depicting blood oxygen level-dependent contrast at rest were also acquired at 3.0T in Massachusetts General Hospital (Boston, USA). The rsfMRI data (N=33) consisted of 150 volumes with the following parameters: number of slices, 42 (interleaved); slice thickness, 3.51 mm; matrix size, 64×64; flip angle, 90°; repetition time (TR), 2400 ms; in-plane resolution, 3.437 mm. In the majority of subjects, DWI was performed using a 3T (Siemens Skyra) scanner with the following parameters: numbers of slices, 160; slice thickness, 5mm; TR, 5500 ms; TE, 99ms; in-plane resolution, 1.375mm. For five of the patients in this cohort 1.5T MRI was used due to medical contraindications for 3T, such as the presence of a pacemaker.

### Image processing

Both structural and functional images were preprocessed using the Configurable Pipeline for the Analysis of Connectomes (CPAC)^45^. First, bias field correction of the anatomical images was performed^46^, followed by brain extraction employing a convolutional neural network^47^. Subsequently, the images were registered to the MNI (Montreal Neurological Institute) anatomical template using non-linear registration (ANTs)^48^. Probability maps for grey matter, white matter and cerebrospinal fluid (CSF) were generated using FSL FAST^49^.

For the functional data, slice timing correction was first performed to account for the interleaved acquisition, while geometrical displacements due to head movement were corrected with rigid registration using the AFNI software (https://afni.nimh.nih.gov/). Brain extraction of the fMRI data was performed using FSL’s Brain Extraction Tool (BET)^50^. The 150 functional images for each patient were then affinely registered to the corresponding T1 image, transformed to the MNI template, and underwent mean intensity normalization. Finally, nuisance signal regression was performed for white matter, CSF and global mean, and the functional time series were band-pass filtered (0.01-0.1Hz) and scrubbed for extreme frame displacement (>3mm). The structural and functional preprocessing steps are summarized in Figure 1.

**Figure 1:**
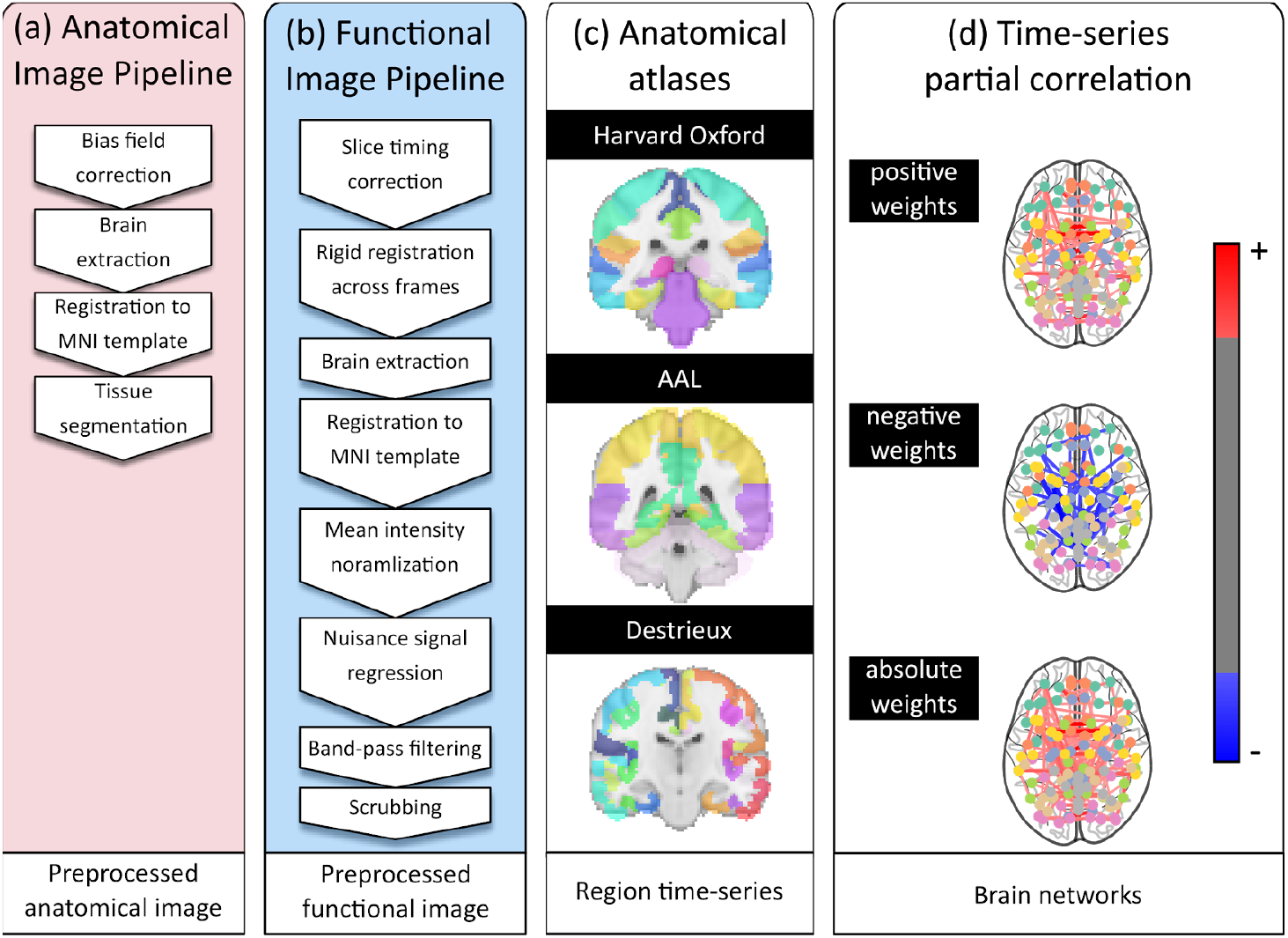
Overview of the processing pipeline. Each patient’s imaging data underwent (a) anatomical processing, (b) functional processing, (c) spatial normalization to one of three atlases (Harvard-Oxford, AAL, Destrieux), and (d) functional connectome creation.

### Rich club region characterization

A study on structural connectivity subdivided the brain into 68 cortical and 14 subcortical regions^51^ and identified those belonging to the RC^8^. These regions are characterized by high connection strength, high betweenness centrality (centrality within a network with respect to its influence on the transfer of information) and low path length (indicator of efficient information transmission). This set comprises 6 bilateral regions, including the *precuneus,* the *superior frontal* and *parietal cortex,* along with subcortical regions including *putamen, hippocampus* and *thalamus* (see Figure 2A). An expert neurologist (M.R.E.), blinded to outcomes, manually identified the number of RC regions affected by the lesion (N_RC_) and outlined the acute infarct lesions on the DWI image.

**Figure 2:**
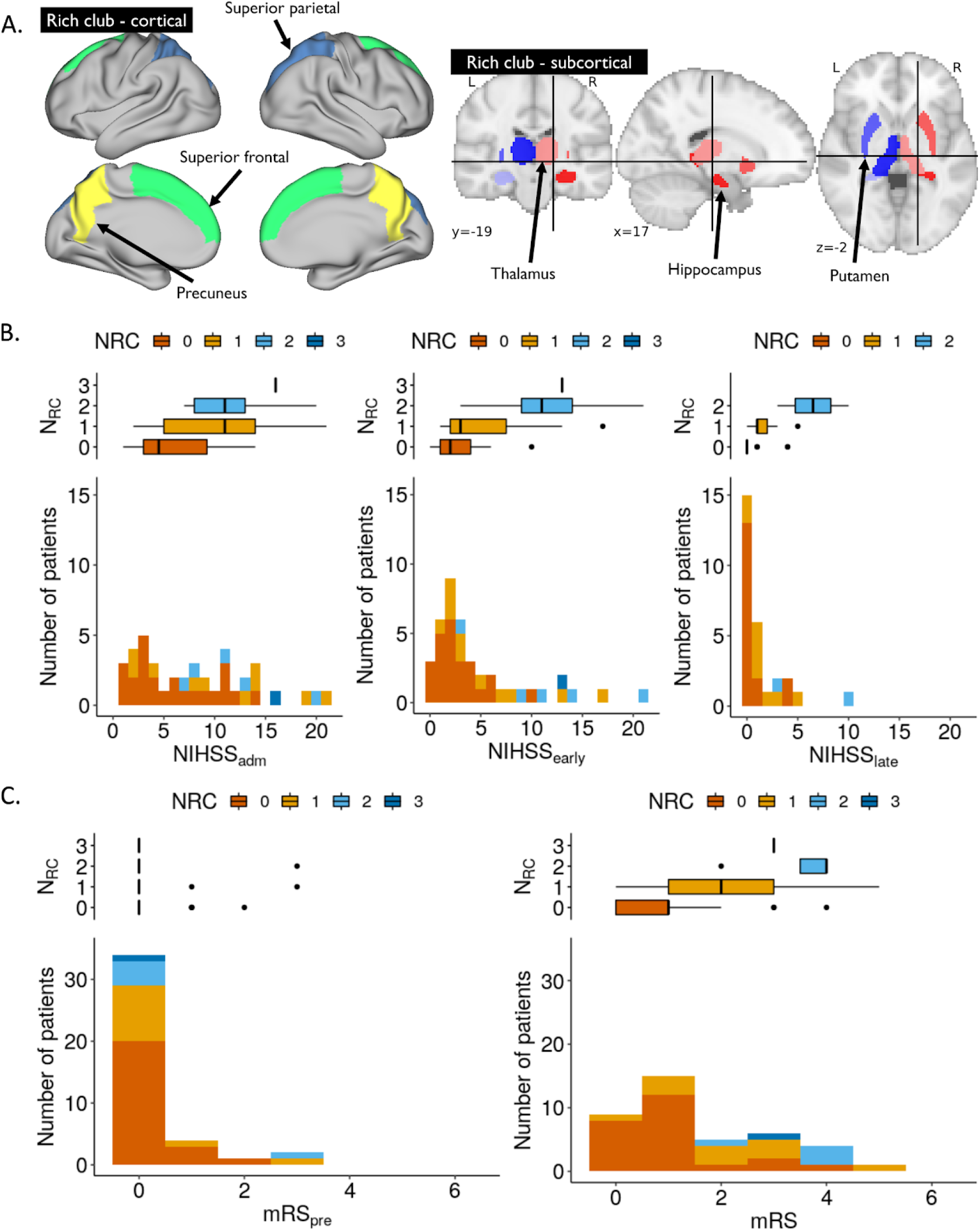
A. Visualisation of cortical (left) and subcortical (right) brain regions comprising the RC backbone. B. Early (2-5 days; N=41; left) and late (90 day; N=28; right) follow-up NIHSS score distribution for all AIS patients, stacked and color-coded by N_RC_. C. Pre-stroke (left) and late (right) mRS assessment for all AIS patients, stacked and color-coded by N_RC_.

### Network analysis of functional connectivity

Three anatomical atlases were used to define the regions of the connectome following preprocessing of the fMRI images, allowing us to explore reproducibility of the findings across different brain parcellations. These included the Destrieux (148 regions)^52^, the Harvard-Oxford^51^, and the AAL atlas (116 regions)^53^. For each atlas and each corresponding regions, mean time series were calculated. Partial correlation between the time series was employed to estimate the strength of the functional connections, yielding a weighted graph representation^54^. Global efficiency of the networks was estimated via the characteristic path length, 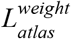, which corresponds to the average path length *l(s)* across all regions *s*^7^, calculated on a given atlas with a given connectivity weight. In this study, we investigated retaining positive, negative and absolute weights of the estimated networks, as there is no consensus with regards to which of these is most discriminative.

### Functional topology and outcome models

We first investigate the association between the number of RC regions affected by stroke (N_RC_) and all outcome measures using the Spearman’s correlation coefficient. Additionally, we assessed the differences in *L* with respect to NIHSS and mRS (‘good’ or ‘poor’ (mRS>2)) based on Pearson’s correlation coefficient and Mann-Whitney-U tests, respectively.

Subsequently, we modeled functional outcome using linear regressions based on age, lesion volume (DWIv), NIHSS at admission (NIHSS_adm_), pre-stroke mRS (mRS_pre_), *L* and N_RC_. First, we performed a univariate analysis between all independent variables and each outcome measure. For multivariate analysis, we defined the *baseline model* based on age, lesion volume (DWIv), and *early outcome measures* (NIHSS_adm_ or mRS_pre_, for outcome models based on NIHSS and mRS, respectively). In the *outcome model,* we further included N_RC_ and *L*. In addition, we considered interactions between N_RC_ and DWIv, due to the fact that larger lesions are likely to affect more RC regions, as well as an interaction term between N_RC_ and *L*, as damage to RC regions have been shown to disturb global network efficiency, and therefore *L*, more significantly compared to other regions^8^. To reduce the statistical burden of the model, we performed backward elimination, where variables with the highest p-value above 0.05 are iteratively eliminated and the model is refit, until only significant terms remain. The outcome models incorporate information related to both structural and functional connectivity, however, as *L* is only available for patients with available fMRI data we remove it after backward elimination to test the model in the larger cohort. The baseline and initial outcome models are given in Table 1.

**Table 1.** Summary of models investigated. Models have the form ‘response ~ terms’, where response is the dependent variable and terms the series of independent variables utilized in the model connected by ‘+’. Interaction terms between independent variables are indicated by ‘:’.

| Model name | Model |
| --- | --- |
| Baseline | Outcome ~ Age + DWIv |
| Initial model | Outcome ~ Age + DWIv + $N_{RC}$ + L + DWIv: $N_{RC}$ + $N_{RC}$:L + early measure |

All models are compared using explained variance with (R^2^_adj_) and without (R^2^) adjustment for the number of independent variables. Furthermore, we report two information criteria, i.e. Akaike information criterion (AIC)^56^ and Bayes information criterion (BIC)^57^, where smaller values correspond to better model fit. BIC, in addition to assessing the model fit, considers a trade-off between model fit and complexity of the model, where more complex models are penalized. All analyses were performed using the computing environment R^55^.

## Results

Forty-four AIS patients were enrolled in this study. Three patients were subsequently excluded because the MRI was not obtained. Of the remaining 41 patients, all had NIHSS at admission and between 2-5 days, as well as mRS recorded. For 28 patients 90-day NIHSS score was also available and fMRI data was collected for 33 patients. Table 2 summarizes the cohort characteristics.

**Table 2.**
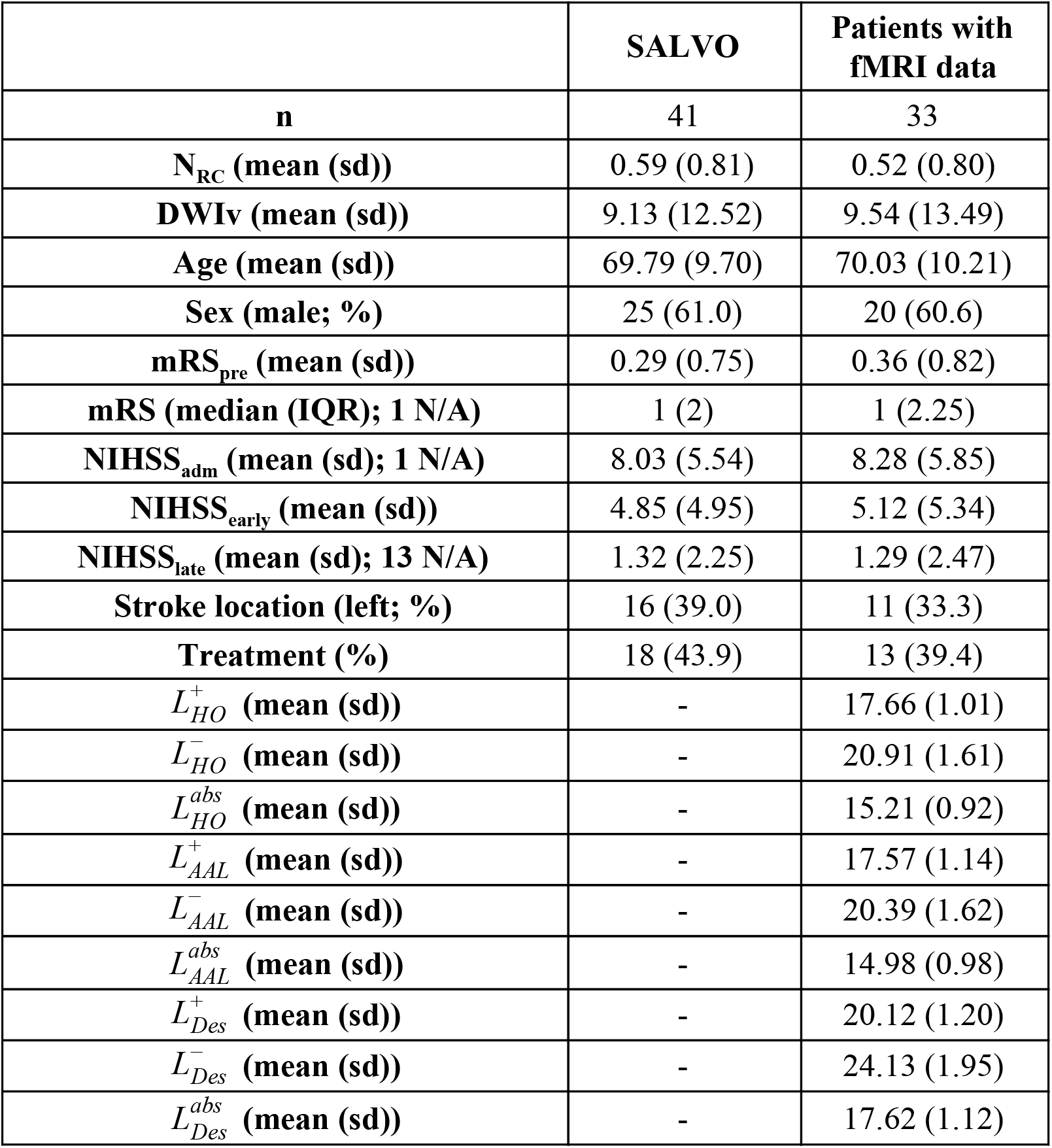
Study cohort characterization. The treatment category includes intravenous tPA or endovascular thrombectomy. Patients with fMRI data available were not significantly different in any of the characteristics (p>0.2). (sd: standard deviation; IQR: inter-quartile range)

### Associations between functional outcome and network topology

Figure 2B characterizes outcomes in our cohort. In our analysis, we observed a positive correlation between all outcome measures and N_RC_ (Spearman’s Rho r = 0.54, 0.58, and 0.58 for NIHSS_early_, NIHSS_late_, and mRS, respectively (all p<0.001); see Figure 2B and C).

After constructing the functional connectivity networks using partial correlation, we explore the *L* for positive weights (mean and standard deviation for all connectomes reported in Table 2; analysis results using all weighting schemes are reported in Table A1). Figure 3A illustrates a positive correlation between the NIHSS_early_ and *L* with regions defined by the Harvard-Oxford, AAL and Destrieux atlases (Pearson’s correlation coefficient r=0.42 (p=0.01), r=0.38 (p=0.03), and r=0.41 (p=0.02), respectively). Additionally, Figure 3B shows the differences in *L* for patients with good and poor outcome at 90 days post-stroke as measured by mRS.

**Figure 3:**
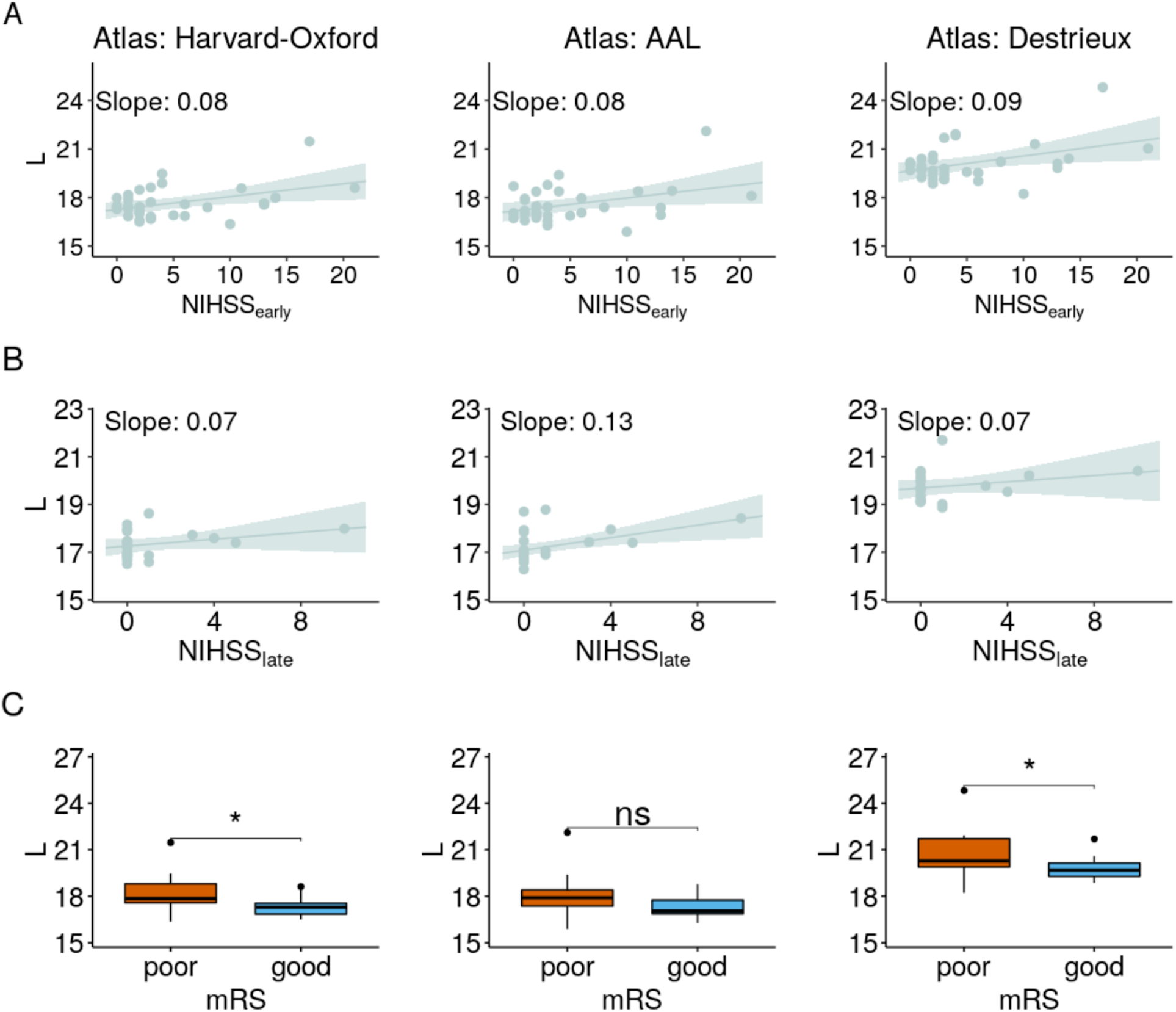
Characteristic path length (L) for networks based on positive correlations and NIHSS_early_ for all atlases. B. *L* for networks based on positive correlations and NIHSS_late_ for all atlases. C. *L* with respect to mRS scores (good outcome (mRS<2). Significance was determined based on Mann-Whitney-U test (*: p≤0.05).

### Outcome model based on connectivity measures

The results of the univariate analysis are shown in Table 3. All factors are significant for all outcome measures, except for mRS_pre_ in case of the 90-day NIHSS score. Results for *L* calculated on the other connectome weighting schemes are summarized in Table A2.

**Table 3.** Summary of univariate analysis for all variables used in the outcome models and *L*, calculated on connectomes retaining positive weights for the Harvard-Oxford (HO), Automated Anatomical Labeling (AAL), and Destrieux (DES) atlases. (*: p<0.05; **: p<0.01; ***:p<0.001)

| | $NIHSS_{early}$ | **p** | $NIHSS_{late}$ | **p** | **mRS** | **p** |
| --- | --- | --- | --- | --- | --- | --- |
| **age** | 0.071±0.011 | *** | 0.019±0.006 | ** | 0.023±0.003 | *** |
| **DWIV** | 0.262±0.057 | *** | 0.129±0.044 | ** | 0.079±0.018 | *** |
| **$N_{RC}$** | 5.450±0.688 | *** | 2.353±0.445 | *** | 1.667±0.236 | *** |
| $L_{HO}^+$ | 0.296±0.051 | *** | 0.075±0.031 | * | 0.092±0.014 | *** |
| $L_{AAL}^+$ | 0.297±0.052 | *** | 0.076±0.031 | * | 0.093±0.014 | *** |
| $L_{Des}^+$ | 0.260±0.045 | *** | 0.066±0.027 | * | 0.081±0.012 | *** |
| **$NIHSS_{adm}$** | 0.560±0.071 | *** | 0.186±0.053 | ** | 0.178±0.020 | *** |
| **$mRS_{pre}$** | 4.923±1.124 | *** | 0.875±0.910 |  | 1.176±0.483 | * |

Utilizing the initial model in the multivariate analysis for NIHSS_early_, and based on the backward elimination, we removed DWIv (p=0.645) and age (p=0.326) from the baseline and DWIv:N_RC_ (p=0.838), DWIv (p=0.864), *L* (p=0.381), and NIHSS_adm_ (p=0.382) from the outcome model. Similarly, we removed age (p=0.967) and NIHSS_adm_ (p=0.176), and DWIv:N_RC_ (p=0.893), *L* (p=0.381), NIHSS_adm_ (p=0.298), and age (p=0.529) from the NIHSS_late_ baseline and outcome model, respectively. Finally, for mRS, we removed mRS_pre_ (p=0.374) and DWIv (p=0.108) from the baseline, as well as DWIv (p=0.944), mRS_pre_ (p=0.786), DWIv:N_RC_ (p=0.581), and age (p=0.391) from the outcome model. The assessment of the different linear regression models after backward elimination are summarized in Table 4.

**Table 4:**
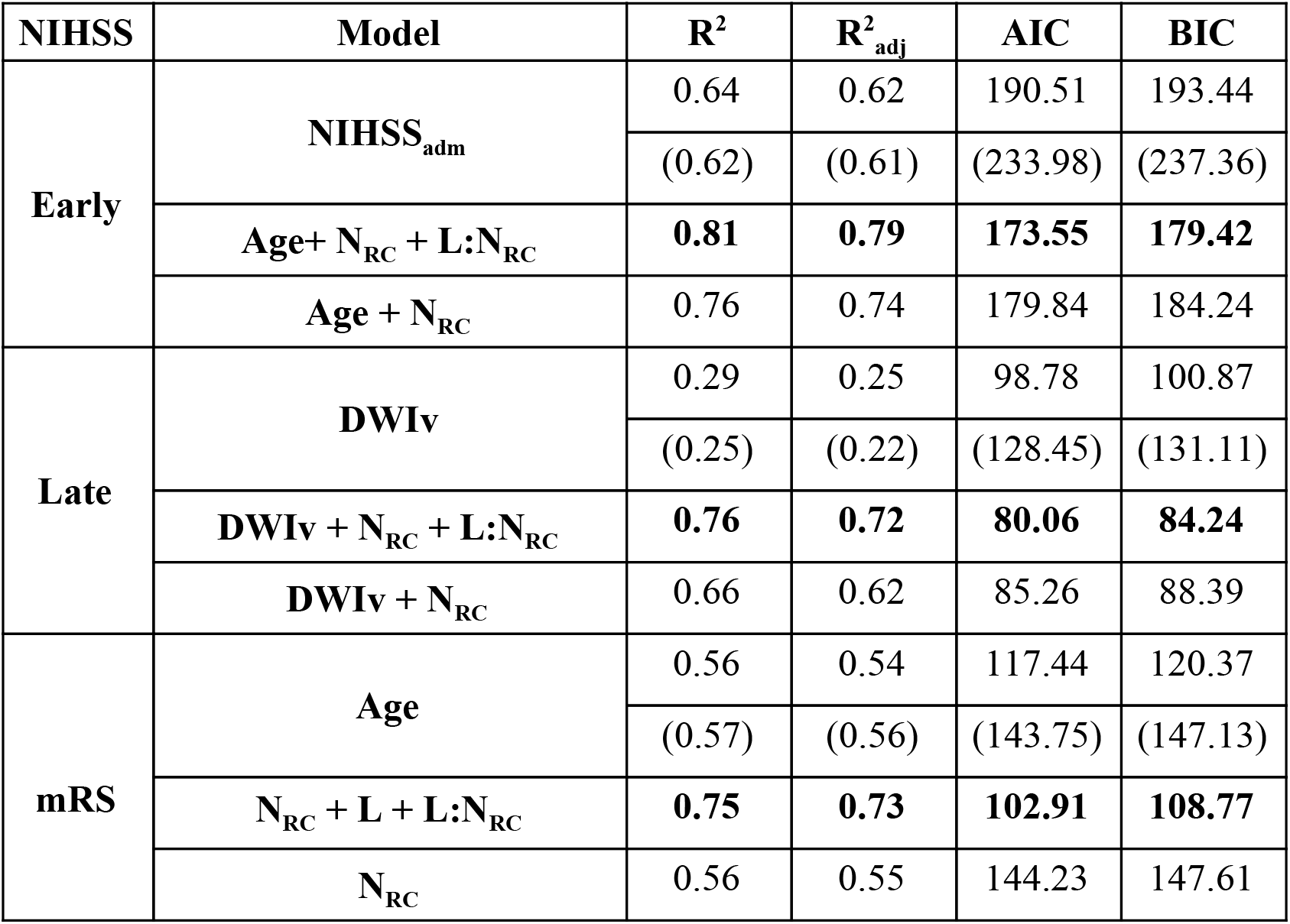
Explained variance for early (2-5 days; top) and late (90-day; bottom) outcome models for subjects with available fMRI data (N=33 and N=21, respectively). Results in parentheses correspond to all available subjects (N=41 and N=28, respectively). Models were compared based on their AIC and BIC values, where the lowest values identify the model that describes the data best (bold).

| NIHSS | Model | R<sup>2</sup> | R<sup>2</sup><sub>adj</sub> | AIC | BIC |
| --- | --- | --- | --- | --- | --- |
| Early | NIHSS<sub>adm</sub> | 0.64 | 0.62 | 190.51 | 193.44 |
|  |  | (0.62) | (0.61) | (233.98) | (237.36) |
|  | Age+ N<sub>RC</sub> + L:N<sub>RC</sub> | **0.81** | **0.79** | **173.55** | **179.42** |
|  | Age + N<sub>RC</sub> | 0.76 | 0.74 | 179.84 | 184.24 |
| Late | DWI<sub>v</sub> | 0.29 | 0.25 | 98.78 | 100.87 |
|  |  | (0.25) | (0.22) | (128.45) | (131.11) |
|  | DWI<sub>v</sub> + N<sub>RC</sub> + L:N<sub>RC</sub> | **0.76** | **0.72** | **80.06** | **84.24** |
|  | DWI<sub>v</sub> + N<sub>RC</sub> | 0.66 | 0.62 | 85.26 | 88.39 |
| mRS | Age | 0.56 | 0.54 | 117.44 | 120.37 |
|  |  | (0.57) | (0.56) | (143.75) | (147.13) |
|  | N<sub>RC</sub> + L + L:N<sub>RC</sub> | **0.75** | **0.73** | **102.91** | **108.77** |
|  | N<sub>RC</sub> | 0.56 | 0.55 | 144.23 | 147.61 |

In all cases, models including connectivity information, i.e. N_RC_ and *L*, resulted in higher explained variance (both unadjusted and adjusted) and lower information criteria (AIC and BIC), compared to all other models. Specifically, including both N_RC_ and *L* resulted in a 1.3-, 2.6-, and 1.3-fold increase in explained variance over the baseline model for NIHSS_early_, NIHSS_late_, and mRS, respectively. Importantly, models after removing *L*, and thereby extending it to the bigger cohort where no fMRI data was available, perform similarly or outperform their corresponding baseline models. Including treatment as a ‘nuisance’ variable in the models did not change performance of any of the models (p>0.2).

## Discussion

Functional outcomes vary significantly in the early and late phases of stroke recovery and are difficult to model at AIS onset. Using the acute stroke lesions visible on the admission DWI, we demonstrated the importance of the integrity of the RC backbone on functional outcomes after AIS, by means of N_RC_, and its association with early and late functional outcome. Additionally, we showed that the topology of functional networks (independent of anatomical atlas) significantly augments the accuracy of outcome models.

Our results align with findings in the literature. Munsch et al.^27^ identified stroke location as an independent determinant of cognitive outcome as measured at 3 months post-stroke. Similar results were demonstrated by Wu et al.^28^, highlighting the importance of joint modelling of DWI volume and topography for stroke outcomes, while our approach also takes network topology into account. Other studies previously have indicated a correlation between network topology and stroke recovery. In their study, Wang et al.^56^ showed that stroke patients exhibit higher network segregation (measured as clustering coefficient^7^) compared to healthy controls and demonstrated an association with restoration of function. Cheng et al.^57^ investigated whole-brain functional network organization in a cohort of 12 stroke patients with motor deficits from 10 days to 3 months post-stroke and showed decreased network integration, corresponding to higher characteristic path length, for patients with right-hemispheric stroke during an ipsilateral finger tapping task. This agrees with our results, where patients with poor outcome showed higher characteristic path length based on their rsfMRI data.

Rich club regions comprise brain areas responsible for distributing a large fraction of the brain’s neural communications. This underpins the importance of these regions for recovery after brain damage, as local disruptions to these central hubs of information flow most likely affect the brain more severely at a global level. The underlying physiological causes of this phenomenon can be explained by (1) the disproportionate impact of pathological attacks on brain hubs on the global efficiency of information processing^58^, (2) the increased vulnerability of these regions to pathogenic factors, due to their topological centrality and high biological cost (manifested by their long-distance neuronal connections), and/or (3) the brain’s inability to compensate for damage or loss of these regions. The outcome model was further improved after introducing a measure of functional network efficiency (L), which directly describes how focal lesions affect the brain network at a global level. After backward elimination in two of the three models only the interaction term between *L* and N_RC_ remained. This suggests that the importance of these regions for functional outcome increases, as efficiency of the brain network decreases (larger *L*. In this case, *L* after stroke might serve as a surrogate measure of *L* before the stroke and future studies are required to disentangle a causal relationship. However, information on *L* before stroke are generally difficult to obtain. Regardless, comparing the associations between *L* and outcome highlights that a more intact and/or efficient network communication in the acute phase of stroke is associated with better outcome, indicated by a lower NIHSS and mRS score. While the mechanisms through which the brain’s functional reorganization facilitates recovery after stroke and the causal relationship between the two variables are yet to be explored, our findings indicate that *L*, estimated from rsfMRI in the acute stroke phase, may be utilized as a determinant of functional recovery. Future studies may also investigate mRS and NIHSS in association with task-specific networks, such as sensorimotor and frontoparietal control systems, to shed light on particular brain regions that are responsible for reorganizations in the connectome.

In general, these results underscore the importance of efficient brain connectivity in functional recovery and resilience to brain damage after ischemic stroke. Our model accounts for some of the most commonly reported confounding factors that are available in the acute setting, i.e. age and lesion volume as measured on the acute DWI image. We found that, although N_RC_ is correlated with DWIv, the volume alone is not enough to explain the early and late outcomes as measured by the NIHSS score. Despite the positive correlation between these two measures (Spearman’s Rho r=0.46), there are both large infarcts that do not affect any RC regions and small infarcts that involve one or two RC regions, demonstrating that a larger DWIv does not equal greater N_RC_. In future work, we aim to investigate whether the incorporation of structural connectivity measures could further improve the prediction accuracy of long-term functional outcome. As suggested by Carter et al.^36^, each behavioral deficit and its variability across stroke patients will likely be explained by a combination of structural variables (e.g. DWIv) and their interaction with measures of structural (e.g. integrity of white matter pathways), and functional connectivity. According to their study, stroke causes a change of ‘functional state’ in the spontaneous brain activity at rest, impacting the brain network during active behavior, further motivating our study in which we investigated resting state activity in isolation and its relation to behavioral deficits and outcome. This is further highlighted by the resulting models after backward elimination, where the models of NIHSS incorporate age or DWIv, as well as measures of structural (N_RC_) and functional (*L*) connectivity.

In this analysis, we saw an increase in explained variance in the outcome model of early NIHSS, where the model using age and connectome information outperform a model using the same measure obtained 2-5 days earlier. While the connectome information remain in the model, age loses its significance and DWIv becomes more important in the late NIHSS assessment. This suggests that age plays an important role in compensating the acute effects of stroke, whereas the effects of structural damage (DWIv) become more important for long-term outcome and recovery. While rsfMRI data is often not available in the hyper-acute stage of stroke, we removed *L* from our models, demonstrating a clinically relevant and easy to assess model. Even without *L*, the presented outcome models demonstrate an increase in explained variance over their corresponding baseline models, except for mRS. However, mRS is the only model not containing either age or DWIv. Re-introducing age into the model, as it was the last parameter removed during backward elimination, results in both age and N_RC_ being significant with explained variance of R^2^=0.66, showing similar improvement compared to the NIHSS based models.

There are limitations that need to be taken into consideration when interpreting these results. First, regional delays have been identified in rsfMRI fluctuations in stroke and cerebrovascular disease patients (hemodynamic lag), which takes place due to vascular occlusion^59^. These delays are measured by time shift analysis of regional BOLD time series with respect to a reference signal. Approaches have been proposed to correct for such lags^60^, however, there is no consensus on how to address this challenge. Importantly, the effect of hemodynamic lag and the subsequent drop in estimated functional connectivity may be an integral part of the observed differences between patients, and may be utilized to determine variations in outcome. Another limitation results from potential registration errors, due to relatively low through-plane resolution of the anatomical scans and the fact that the lesions were not masked out when registering the anatomical scans to the MNI template. While registration errors increase noise in the analysis, it is unlikely that this will cause a systematic error in our cohort of patients with right- (N=22)) and left-hemispheric (N=11) strokes. Moreover, by using three atlases to investigate the functional network topology and demonstrating consistent trends, our results are less prone to systematic errors due to misalignment of boundaries between regions. In this study, a subset of patients (N=5) had contraindication for 3T MRI acquisition and subsequently underwent 1.5T imaging. However, studies suggest that there are no significant differences in the assessment of infarct lesion volume between 1.5T and 3T systems in the hyperacute stage^61^. Generally, this study presents a proof-of-concept, due to the relatively small sample size. In particular, only few subjects showed poor outcome, which can affect model fit, as these might be considered ‘outliers’. While the assumptions of the linear models, i.e. mean residual equals 0, no correlation between residuals and dependent variables, positive variability, homoscedasticity, and no multicollinearity (defined as variance inflation factors < 2), were fulfilled, we observed a quantile-quantile plot corresponding to a heavy-tailed distribution of the standardized residuals. However, after excluding subjects with ‘extreme’ outcome values (NIHSS_early_>15, NIHSS_late_>9, and mRS of 5), thereby improving model fit, the results remained consistent and demonstrated an increase in explained variance. Moreover, mRS is usually modeled using ordinal regressions, requiring larger datasets than was available in this study. Considering the general agreement with outcome models using NIHSS in terms of factors that are retained in the analysis, however, we do not expect a significant change in retained factors, as using linear regression is more likely to introduce more noise in the data.

Among the strengths of this study was the thoroughly ascertained and well-characterized hospital-based dataset of patients with AIS and consecutive assessments of functional post-stroke outcomes. These included consecutive outcome measurements of NIHSS, which is a fine-grained measure for quantifying the impairment of stroke. Importantly, in combining N_RC_ and topological information from functional connectivity profiles of patients, we were also able to interrogate effects on both the structural (N_RC_) and functional (*L*) brain networks, combining them into a single, intuitive outcome model.

## Summary

In conclusion, this is the first study exploring functional network and RC topology of brain connectivity in AIS patients, as well as their association with early and late post-stroke outcomes. Our findings highlight the impact of stroke location on functional recovery, as well as the importance of structural connectivity hubs and functional integration for efficient information transmission. The proposed model yields a 1.3-2.6-fold improvement in explained variance over the baseline model, improving our understanding of how stroke affects functional brain organization in the acute setting.

### Source of Funding

This work was funded by NIH Grant 5R01NS082285. Sofia Ira Ktena was supported by the EPSRC Centre for Doctoral Training in High Performance Embedded and Distributed Systems (HiPEDS, Grant Reference EP/L016796/1) and an EMBO short-term fellowship (Reference 7284). Markus D. Schirmer was supported by the European Union’s Horizon 2020 research and innovation programme under the Marie Sklodowska-Curie grant agreement No 753896.

Disclosures

---

**Table A1.** Summary of results for functional connectomes created retaining positive, negative, and absolute connectivity profiles for the Harvard-Oxford (HO), Automated Anatomical Labeling (AAL), and Destrieux (DES) atlases. Path-lengths *L* were calculated for each atlas and each connectome combination and assessed with respect to their Pearson’s correlation coefficient with early and late outcome, measured by NIHSS. Additionally, the explained variance with and without adjustment for number of independent variables (R^2^ and R^2^_adj_), and the Bayes information criterion (BIC) are reported for the outcome models in Table 4.

| Connectome | | Pearson's Correlation Coefficient ($L \sim \text{NIHSS}$) | | L(mRS)<br>(mean (std)) | | | Outcome model after backward elimination | | | | | | | | |
| --- | --- | --- | --- | --- | --- | --- | --- | --- | --- | --- | --- | --- | --- | --- | --- |
|  |  |  |  |  |  |  | NIHSS<sub>early</sub> |  |  | NIHSS<sub>late</sub> |  |  | mRS |  |  |
| Atlas | Weights | Early | Late | good | poor | p | $R^2$ | $R^2_{adj}$ | BIC | $R^2$ | $R^2_{adj}$ | BIC | $R^2$ | $R^2_{adj}$ | BIC |
| HO | positive | 0.42* | 0.32 | 17.3 (0.6) | 18.3 (1.4) | * | 0.82 | 0.80 | 178.05 | 0.67 | 0.61 | 87.20 | 0.75 | 0.72 | 109.68 |
|  | negative | 0.26 | -0.03 | 20.5 (1.0) | 21.8 (2.3) | - | 0.81 | 0.79 | 179.79 | 0.67 | 0.61 | 87.47 | 0.73 | 0.70 | 111.86 |
|  | absolute | 0.37* | 0.15 | 14.9 (0.6) | 15.8 (1.3) | * | 0.82 | 0.80 | 178.54 | 0.66 | 0.60 | 87.69 | 0.74 | 0.71 | 110.56 |
| AAL | positive | 0.38* | 0.42 | 17.3 (0.7) | 18.2 (1.7) | - | 0.81 | 0.79 | 179.42 | 0.75 | 0.71 | 81.53 | 0.75 | 0.73 | 108.77 |
|  | negative | 0.19 | -0.08 | 20.1 (1.2) | 21.1 (2.3) | - | 0.79 | 0.76 | 183.33 | 0.66 | 0.60 | 87.67 | 0.73 | 0.70 | 111.57 |
|  | absolute | 0.30 | 0.16 | 14.8 (0.6) | 15.5 (1.5) | - | 0.80 | 0.78 | 181.33 | 0.68 | 0.62 | 86.59 | 0.75 | 0.72 | 109.58 |
| DES | positive | 0.41* | 0.25 | 19.8 (0.7) | 20.8 (1.8) | * | 0.81 | 0.79 | 179.57 | 0.66 | 0.60 | 87.61 | 0.74 | 0.71 | 110.20 |
|  | negative | 0.32 | -0.07 | 23.6 (1.2) | 25.2 (2.8) | * | 0.80 | 0.78 | 181.01 | 0.68 | 0.63 | 86.35 | 0.73 | 0.70 | 111.54 |
|  | absolute | 0.39* | 0.12 | 17.3 (0.6) | 18.3 (1.7) | * | 0.81 | 0.79 | 179.35 | 0.67 | 0.61 | 87.56 | 0.74 | 0.71 | 110.59 |

**Table A2.** Summary of univariate analysis results for all variables used in the outcome models.

|  | NIHSS<sub>early</sub> | p | NIHSS<sub>late</sub> | p | mRS | p |
| --- | --- | --- | --- | --- | --- | --- |
| **age** | 0.071±0.011 | *** | 0.019±0.006 | ** | 0.023±0.003 | *** |
| **DWIV** | 0.262±0.057 | *** | 0.129±0.044 | ** | 0.079±0.018 | *** |
| **N<sub>RC</sub>** | 5.450±0.688 | *** | 2.353±0.445 | *** | 1.667±0.236 | *** |
| **NIHSS<sub>adm</sub>** | 0.560±0.071 | *** | 0.186±0.053 | ** | 0.178±0.020 | *** |
| **mRS<sub>pre</sub>** | 4.923±1.124 | *** | 0.875±0.910 |  | 1.176±0.483 | * |
| $L_{HO}^+$ | 0.296±0.051 | *** | 0.075±0.031 | * | 0.092±0.014 | *** |
| $L_{HO}^-$ | 0.248±0.044 | *** | 0.063±0.027 | * | 0.078±0.012 | *** |
| $L_{HO}^{abs}$ | 0.343±0.060 | *** | 0.087±0.036 | * | 0.107±0.016 | *** |
| $L_{AAL}^+$ | 0.297±0.052 | *** | 0.076±0.031 | * | 0.093±0.014 | *** |
| $L_{AAL}^-$ | 0.253±0.045 | *** | 0.063±0.027 | * | 0.079±0.012 | *** |
| $L_{AAL}^{abs}$ | 0.347±0.061 | *** | 0.088±0.036 | * | 0.108±0.017 | *** |
| $L_{Des}^+$ | 0.260±0.045 | *** | 0.066±0.027 | * | 0.081±0.012 | *** |
| $L_{Des}^-$ | 0.216±0.038 | *** | 0.054±0.023 | * | 0.068±0.010 | *** |
| $L_{Des}^{abs}$ | 0.297±0.051 | *** | 0.075±0.031 | * | 0.093±0.014 | *** |

